# Mobility can promote the evolution of cooperation via emergent self-assortment dynamics

**DOI:** 10.1101/175638

**Authors:** Jaideep Joshi, Iain D Couzin, Simon A Levin, Vishwesha Guttal

## Abstract

The evolution of costly cooperation, where cooperators pay a personal cost to benefit others, requires that cooperators interact more frequently with other cooperators. This condition, called positive assortment, is known to occur in spatially-structured viscous populations, where individuals typically have low mobility and limited dispersal. However many social organisms across taxa, from cells and bacteria, to birds, fish and ungulates, are mobile, and live in populations with considerable inter-group mixing. In the absence of information regarding others’ traits or conditional strategies, such mixing may inhibit assortment and limit the potential for cooperation to evolve. Here we employ spatially-explicit individual-based evolutionary simulations to incorporate costs and benefits of two coevolving costly traits: cooperative and local cohesive tendencies. We demonstrate that, despite possessing no information about others’ traits or payoffs, mobility (via self-propulsion or environmental forcing) facilitates assortment of cooperators via a dynamically evolving difference in the cohesive tendencies of cooperators and defectors. We show analytically that this assortment can also be viewed in a multilevel selection framework, where selection for cooperation among emergent groups can overcome selection against cooperators within the groups. As a result of these dynamics, we find an oscillatory pattern of cooperation and defection that maintains cooperation even in the absence of well known mechanisms such as kin interactions, reciprocity, local dispersal or conditional strategies that require information on others’ strategies or payoffs. Our results offer insights into differential adhesion based mechanisms for positive assortment and reveal the possibility of cooperative aggregations in dynamic fission-fusion populations.

**Author Summary:** Cooperation among animals is ubiquitous. In a cooperative interaction, the cooperator confers a benefit to its partner at a personal cost. How does natural selection favour such a costly behaviour? Classical theories argue that cooperative interactions among genetic relatives, reciprocal cooperators, or among individuals within groups in viscous population structures are necessary to maintain cooperation. However, many organisms are mobile, and live in dynamic (fission-fusion) groups that constantly merge and split. In such populations, the above mechanisms may be inadequate to explain cooperation. Here, we develop a minimal model that explicitly accounts for mobility and cohesion among organisms. We find that mobility can support cooperation via emergent dynamic groups, even in the absence of previously known mechanisms. Our results may offer insights into the evolution of cooperation in animals that live in fission fusion groups, such as birds, fish or mammals, or microbes living in turbulent media, such as in oceans or in the bloodstreams of animal hosts.

## Introduction

In a cooperative interaction, the actor’s actions benefit others. In some cases, cooperators bear a fitness cost. Such costly cooperative, or ‘non-selfish’, traits are typically selected against in well mixed populations. However, cooperation can evolve if cooperators are more likely to interact with other cooperators, i.e. if there is ‘positive assortment’ of the cooperative allele [1, 2]. There are several well known mechanisms that lead to positive assortment. Preferential interactions among close genetic relatives can create assortment of cooperators by common descent [3]. Alternatively, tag-based interactions, where a phenotypic tag (also called ‘greenbeard’) mediates preferential interactions among cooperators [4–6], can also create assortment. In the absence of abilities to recognise other individuals or their traits, spatial structures arising from local dispersal of offspring and low mobility can facilitate assortment by creating local clusters of cooperators [7–12]. Elucidating various mechanisms of positive assortment has been a fundamental question in evolutionary biology.

Populations of many social organisms, from cells and bacteria to many species of fish and birds, consist of dynamic groups that frequently merge and split. In such populations, also called ‘fission-fusion populations’ [13], mobility (low ‘population viscosity’) causes considerable mixing among groups. This reduces relatedness among members of groups [14, 15], limiting interactions among close genetic relatives. Therefore, the potential for assortment based on common descent would seemingly be rare. Mobility also leads to paucity of repeated interactions between individuals, potentially limiting reciprocal cooperation [16]. Spatial structure arising from the presence of multiple groups in such populations could, in principle lead to assortment of cooperators [1,.17]. However, positive assortment cannot happen in randomly formed groups [1].Rather, it is typically expected that individuals remain in their natal groups, at least over one generation, to enhance assortment [8,.9]. Groups members can also adopt conditional strategies, where they can respond to others’ behavioural or group traits to avoid non-cooperators [18–21] and to prefer interactions with cooperators. These scenarios are unlikely in fission-fusion populations where groups frequently merge and split. Moreover, previous evolutionary theory has demonstrated that even slight mixing among groups would be expected to abolish any realistic possibility of assortment [1, 22]. Consequently, it is typically thought that the selfish interest of individuals takes precedence over group-level benefits in fission-fusion populations [23].

However, organisms living in dynamic social groups do exhibit various cooperative behaviours, such as predator inspection [24, 25], collective mobbing of predators [26, 27] and sentinel behaviours [28] that have the potential to put cooperators at risk. Furthermore, cooperation is increasingly becoming evident in the cellular domain, such as cooperative regulation of growth rates [29], and altruistic suicide by cells to benefit neighbouring cells [30, 31]. Such microbial populations are also often mobile, either through self-propulsion or via environmental forcing, forming fission-fusion groups [32]. Classical theory shows that sufficiently high mobility will undercut selection for cooperation [14, 33, 34]. One mechanism that generates spontaneous sorting of phenotypes in mobile social groups is differential cohesion between cooperators and defectors [35–37]. However, defectors could evolve to match the cohesion of cooperators, thus eliminating differential cohesion driven spatial sorting of cooperators. Here, we demonstrate that explicitly accounting for the spatio-temporal nature of movement and coevolutionary dynamics between cooperation and cohesion can resolve this potential paradox, revealing a dynamic and emergent interplay between cooperative trait evolution and assorted spatial structure within populations.

## Model and Methods

We consider self-interested (selfish) individuals within a continuous two-dimensional spatial environment. To account for different ways in which organismal motility could arise, we consider two scenarios. The first is where individuals are capable of mobility via self-propulsion, as is characteristic of many animal populations (such as schooling fish, flocking birds, herding ungulates). The second (at an opposite extreme), is where individuals have no capacity for self-propulsion, but are moved by the flow of the medium in which they live, representing organisms in turbulent media (such as microbes). These two cases are referred to as “active”and “passive” systems, respectively (see S1 Appendix). We measure individual mobility in these two systems by the individual propulsion speed or the average fluid speed of the medium, respectively. Our active system model is based on previous computational frameworks that investigated the evolution of collective movement strategies under ecological scenarios like migration or cannibalism [38, 39]. Our passive system model is based on the work of Torney et. al [40], which investigated collective search behaviours in dynamic environments. Here, we employ these model frameworks but with different individual payoff structures to study the coevolutionary dynamics of cooperation and cohesion.

### Individual-based evolutionary model

In both active and passive system models, we assume that individuals have two evolvable behavioural traits. The first trait is a local tendency to be cohesive with neighbours, denoted by a continuous variable *ω*_*s,i*_*∈* [ 0, ∞) (e.g. via social interactions among animals, or regulation of cellular adhesion among microbes). Cohesive interactions cause formation of groups; we define any two individuals to be in the same group if the distance between them is less than a threshold denoted by *R*_*g*_. Depending on the strength of cohesive tendencies among individuals, populations can exhibit a variety of dynamic group formations, as found in social organisms (Supplementary Video 1). These can range from large cohesive groups (where individuals exhibit high cohesion, or large *ω*_*s,i*_) to largely solitary motion (when individuals exhibit no cohesion, *ω*_*s,i*_ = 0). We assume that individuals pay a fitness cost proportional to the strength of their cohesive tendencies (*c*_*s*_(1 + *ω*_*s,i*_)^2^ for the active system and 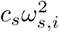 for the passive system; see S1 Appendix for details). This could represent direct costs associated with animals scanning their local area for neighbours, or production of sticky extracellular matrix by cells [41]; and indirect costs such as sharing resources or increased susceptibility to diseases associated with group formation [42].

We consider a second, binary trait that represents strategies of cooperation or defection by individuals. In a pairwise interaction modelled as a prisoner’s dilemma, cooperation results in a personal cost *c* to the donor but offers a benefit *b* to the recipient (*b > c >* 0). Generalizing this to any spontaneously formed group *g* containing *n*_*g*_ (*≥* 2) individuals of which *k*_*g*_ are cooperators, the pay-off to cooperators is (*k*_*g*_ - 1)*b/*(*n*_*g*_ - 1) - *c*, and that to defectors is *k*_*g*_*b/*(*n*_*g*_ - 1). Solitary cooperators get a payoff of -*c* and solitary defectors get zero payoff. Consequently, irrespective of group size and composition, defectors in the group always receive a higher payoff than cooperators. This pay-off structure excludes self-benefits and is also called strong altruism, which evolves only in a very narrow range of conditions [43].

The fitness of each individual is determined from a baseline payoff plus the payoffs associated with cooperative interactions minus the costs of cohesiveness. We assume non-overlapping generations. Therefore, individuals reproduce with a probability proportional to their relative fitness in the population, and then die. Offspring inherit both traits with a small probability of mutation and are dispersed to random locations in space. Such random dispersal makes the average relatedness of individuals within any local neighbourhood equal to the population average, as has been reported in fission-fusion societies [15]. Thus, we have deliberately assumed a simple scenario involving non-repeated cooperative games among non-kin where individuals cannot assess others’ behavioural traits or payoffs of interactions. Thus, mechanisms of cooperation such as kin-interactions, local dispersal and conditional strategies are ruled out. Please see S1 Appendix for more details of the model and other pay-off matrices.

We define population-level trait values of cooperation and cohesion (denoted by *p* and *ω*_*s*_) as averages of the individual trait values (*ω*_*c,i*_ and *ω*_*s,i*_) over the population for a given generation. We then define the ‘evolved cooperative tendency’ (denoted by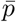), and ‘evolved cohesive tendency’ (denoted by 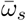) as an average of the respective population-level traits over 1500 generations, further averaged across ten replicate populations (see S1 Appendix). We examine evolved cooperative and cohesive tendencies as compared to baseline cases. The baseline for cohesive tendencies is obtained by setting the proportion of cooperators to zero (i.e. only the cohesive tendencies evolve). There are two plausible baselines for cooperative tendencies, both representing well-mixed systems in which cooperation is not expected to evolve if the population size is infinity: 1) Solitary baseline-this is obtained assuming that the cohesive tendency for all individuals is zero (*ω*_*s,i*_ = 0); therefore, cooperative interactions occur due to chance encounters alone. 2) Single group baseline-this is obtained assuming that the cost of cohesion is zero, which results in all individuals to evolve large cohesive tendencies (i.e. effectively *ω*_*s,i*_*→ ∞*) and form a single cohesive group. The interactions would then resemble those in a well-mixed population. Although other baseline scenarios that represent well-mixed systems can, in principle, be constructed, e.g. all individuals have an intermediate value of cohesive tendencies, one of our goals is to investigate if the coevolutionary dynamics between cooperation and cohesion may indeed select for an intermediate value of cohesive tendencies. Therefore, we avoid intermediate values of *ω*_*s,i*_ as baseline scenarios.

## Results

### Evolution of cooperation and fission-fusion dynamics

We find that in both active and passive systems, mobility facilitates the evolution of cooperation together with local cohesive tendencies, over a range of values spanning several orders of magnitude (Fig 1a-d). Consequently, the evolved populations exhibit merge and split group dynamics, and at the same time, there exists coexistence of cooperators and defectors (Fig 1a-b and Supplementary Videos 1-2). The evolved proportion of cooperators is much higher when both the cohesive and cooperative traits coevolve, as opposed to when only the cooperative trait, but not the cohesive trait, can evolve (black smooth line versus the broken black line in Fig 1c-d; also see S3 Appendix). Likewise, the evolved cohesive tendency is higher in the coevolutionary scenario when compared to cases where only the cohesive tendency, but not the cooperative trait, evolves (blue smooth line versus the broken blue line in Fig 1c-d; also see S3 Appendix).

**Fig 1.**
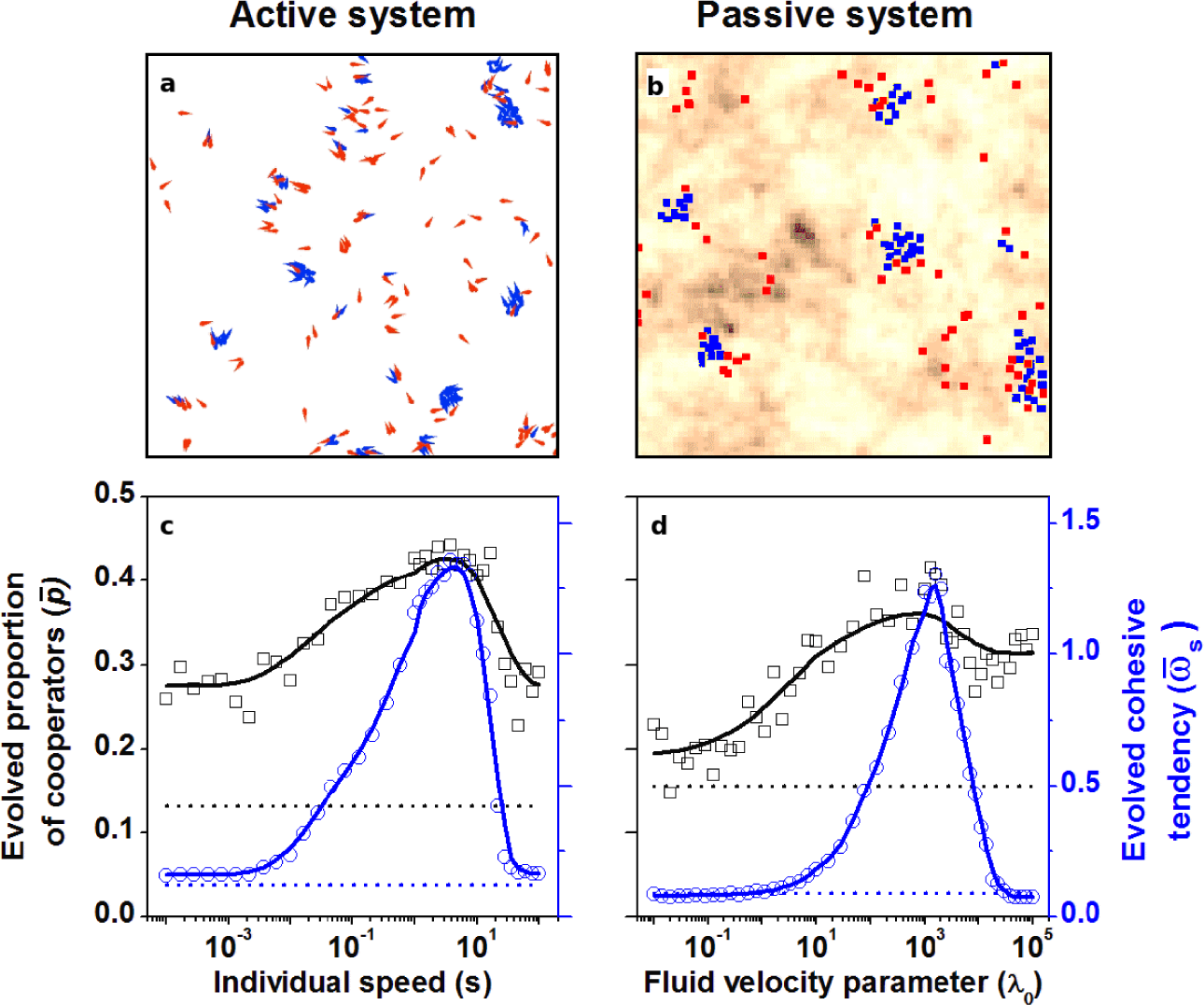
Mobility promotes the evolution of cooperation together with a process for assortment. Representative snapshots of evolved self-sorted group structures containing cooperators (blue) and defectors (red) in (a) the active system where individuals are self-propelled particles, and the passive system where individuals follow the flow of a dynamic medium (background colours represent the potential field of the fluid, with average fluid speed of one unit.) We have exaggerated parameter values to make the assortment visually apparent (see Supplementary Video 2; Fig 3c). (c) and (d) show the evolved proportion of cooperators (*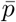*, black squares and smoothed line) and evolved cohesive tendency (*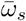*, blue circles and smoothed line), for active and passive systems, respectively, for a range of mobility parameters spread over six orders of magnitude. Both evolved traits are minimal at very low and high mobilities because they do not allow formation of group structures. The black broken line indicates a baseline of evolved cooperation when cohesive tendency is fixed at zero, whereas the blue broken line indicates the baseline of cohesive tendency in the absence of cooperation (S3 Appendix). Simulation parameters: Turbulence parameter *μ* = 0.3, cost of cooperation *c* = 0.1, cost of cohesion *c*_*s*_ = 2. Other parameters are as in Table 1 (see S1 Appendix).

In both active and passive systems, the evolved proportion of cooperators and their associated cohesive tendencies reduce as a function of costs of cooperation, as intuitively expected (Fig 2a,b;e,f). As cost of cohesion increases, we find that cooperators increase in the population (Fig 2 c,d) but this effect is small for the passive system. However, the overall cohesive tendencies decrease with cost of cohesion (Fig 2g,h). Furthermore, the coevolutionary scenario leads to increased levels of cooperation and cohesion in comparison to a case where only the cooperative trait (solitary and single group baselines in Fig 2a, b), or the cohesive trait (‘No coop’ baseline in Fig 2g, h) is allowed to evolve. We also observe that the coevolutionary dynamics of cooperation and cohesion do not reach a steady state, and instead fluctuate over time (Fig 3a for active system, S3 Figure-a for passive system). Except when the costs of cohesion or cooperation are high, evolved populations exhibit fission-fusion dynamics.

**Fig 2.**
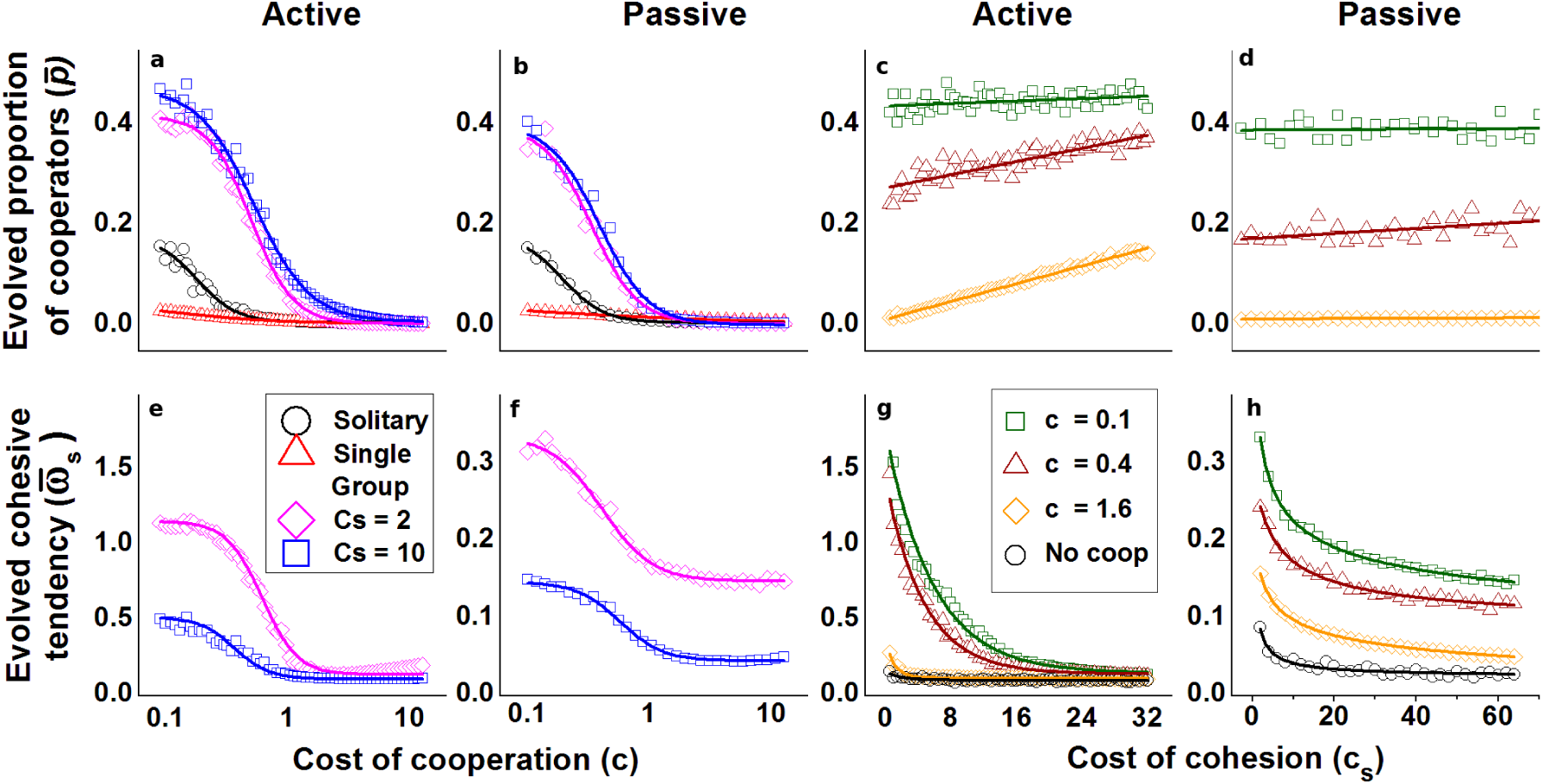
The coevolved proportion of cooperators and average cohesive tendency are higher than baseline scenarios with no coevolution. The evolved proportion of cooperators 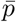 reduces with cost of cooperation (a,b), and increases with increasing cost of cohesion (c,d). The coevolved cohesive tendency 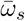 reduces as a function of cost of cooperation (e,f) and cost of cohesion (g,h). Legends and baselines: (i) ‘Solitary’ is a baseline for *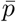*of a well mixed system where individuals walk randomly (*ω*_*s,i*_ = 0 ∀ *i*) and group formation happens by accidental encounters alone. (ii) ‘Single group’ is another baseline for 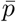, also representing a well mixed system, where all individuals have a strong cohesive tendency and form a single large group. These two baselines are applicable to (a,b).‘No coop’ is a baseline for 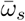in the absence of cooperation. This baseline is applicable to (g,h). (see S3 Appendix for details of baselines). Parameters: Movement speed *s* = 1 (active case), *μ* = 0.1, fluid velocity parameter *λ*_0_ = 1 (passive case), benefit of cooepration *b* = 100. Other parameters as in Table 1 (see S1 Appendix).

We find that our results are robust to a number of modifications to model parameters and specifications, such as the criterion we used for defining a group, the system size of the simulations, specific rules of interactions among neighbours, and the turbulence intensity (S1 Appendix). In addition, we also modified the payoff structure of cooperation: In the standard multiplayer prisoner’s dilemma that we have studied so far, benefits of cooperation are shared equally among all group members (except the cooperator who contributed that benefit). The cost of cooperation, however, is not shared. This is a reasonable representation of many biological systems, especially in the microbial world [30, 44]. However, in scenarios like alarm calling, where an individual calls to warn others of an approaching predator, the benefits of the call may be available fully to all nearby individuals even if a single individual produces a call. The payoff is therefore not divided among group members. To account for such scenarios, we considered payoff structures where the benefits and the costs may or may not depend on the group size. We find that our results are robust to such variations (S5 Appendix).

### Emergence of assortment

We derive the condition for increase in cooperators for any given population structure in our model using first principles, i.e. by computing the expected number of offspring of cooperators and defectors in the next generation (S3 Appendix). Since our model assumes synchronous global reproduction and random dispersal, we obtain the same condition starting with the Price Equation (see S4 Appendix). We then derive the assortment of the cooperative genotype.

The condition for increase in cooperators is *rb > c*, where

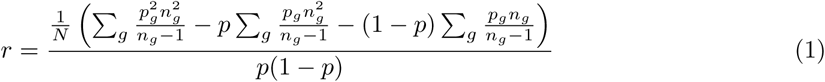

where *p* is the total proportion of cooperators (i.e. the average cooperative tendency of the population at a given generation), *p*_*g*_ is the proportion of cooperators in group *g*, and *n*_*g*_ is the size of group *g*. The summations are performed over groups of size greater than one.

The variable *r* in expression 1 is identical to the assortment of the cooperative genotype, defined as the coefficient of regression of the actor’s genotype on the genotype of its co-players [1, 2]:

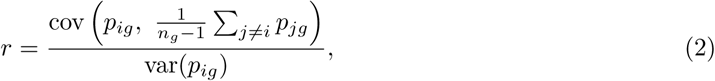

where, *p*_*ig*_ is the indicator variable that individual *i* in group *g* is a cooperator.

We compute *r* from the emergent population and group structure in our simulations, and find that cooperators increase in frequency when *rb >c*, as per the theoretical prediction (Fig 3b; S3 Figure-b). We emphasize that we began our simulations with all individuals being defectors and having no cohesive tendencies. The evolved populations where individuals have local cohesive interactions led to group structures with positive assortment. In other words, assortment is not imposed by our simulations. Rather, it is a consequence of coevolutionary dynamics of cooperation and cohesive interactions.

**Fig 3.**
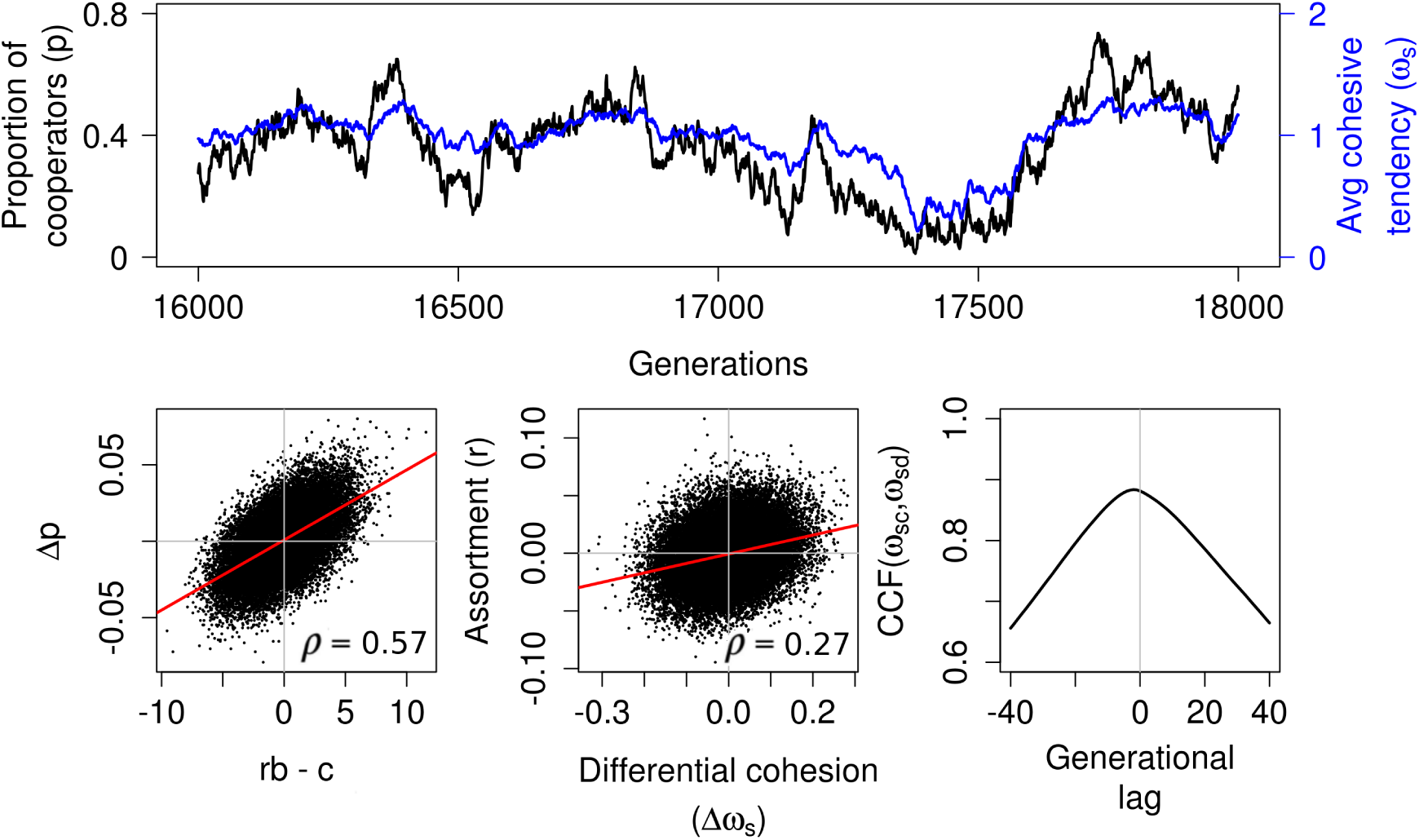
The coevolutionary dynamics and emergent assortment driven by evolving differential cohesive tendencies. (a) Temporal variations in varying *p* (black) and *ω*_*s*_ (blue) suggesting non-equilibrium coevolution dynamics between cooperative and cohesive strategies. (b) The change in proportion of cooperators (Δ*p*) per generation is proportional to *rb - c*, consistent with theoretical predictions. (c) Positive assortment (*r >* 0) results from emergent differential cohesive tendency (Δ*ω*_*s*_ = *ω*_*sc*_ - *ω*_*sd*_ *>* 0) of cooperators and defectors. In b-c, actual data points from the simulation are plotted along with the regression line and correlation coefficient (*ρ*). Both correlations are significant with p-values < 2.2 × 10^-16^ (d) Cross correlation function (CCF) between average cohesive tendencies of cooperators (*ω*_*sc*_) and defectors (*ω*_*sd*_) shows that defectors lag cooperators on average by about 1 generation, suggesting an arms race like dynamics. Parameters: *c* = 0.1, *b* = 100, *c*_*s*_ = 2. Other parameters as in Table 1 (see S1 Appendix).

### Evolved differences in cohesive tendencies create assortment

We now address the question: What are the proximate and ultimate mechanisms for the emergence of a group structure with positive assortment? The answer lies in the coevolutionary dynamics, and spontaneous “self-sorting”within populations [35, 36]. The average cohesive tendency of cooperators within the population (*ω*_*sc*_) differs from that of defectors (*ω*_*sd*_) in every generation, even though the average difference across generations is zero. This variation in social cohesion among individuals results in group-level assortment of individuals based on the cohesive trait (Fig 3c), as has previously been hypothesized for animal and cellular groups [35–37, 45]. Cooperators benefit from positive assortment when their cohesive tendency is higher than that of defectors (i.e. Δ*ω*_*s*_ = *ω*_*sc*_ - *ω*_*sd*_ *>* 0; Fig 3c; S3 Figure-c). By contrast, defectors benefit by having the same cohesive tendency as cooperators because of increased likelihood of joining cooperator groups. Our analysis of the time series of the population cohesive tendency shows that *ω*_*sd*_ lags behind *ω*_*sc*_ by one generation (Fig 3d; S3 Figure-d). This means that cooperators evolve higher *ω*_*sc*_ than defectors to achieve positive assortment, but defectors evolve to match their *ω*_*sd*_ to that of cooperators. This resembles an arms race between them over the costly cohesive trait [46], which continues until the cost of cohesion can no longer be offset by the benefits of assortment.

### Analytical model for coevolutionary arms race dynamics

#### Simplified model and analytical approximation

To further investigate the arms race dynamics, we develop an analytical model for a simplified version of our system. Here too, we consider two evolvable traits: (i) a binary cooperative trait *ω*_*c,i*_ as before, i.e. cooperation or defection, and (ii) a binary cohesive trait, where individuals are either highly cohesive (*ω*_*s,i*_ = 1) or non-cohesive (*ω*_*s,i*_ = 0), in contrast to the continuous valued trait used earlier. This simplification to a binary trait is motivated by the results of the previous section which showed that cooperators and defectors evolve differential cohesive tendencies. Despite the above simplification, numerical simulations reveal that this model can capture various possible movement strategies, from solitary movement (when all individuals are non-cohesive) to fission-fusion group dynamics (when all individuals are cohesive). Besides analytical tractability, a key advantage of this binary trait model is that it amplifies the payoff differences between different types of individuals. As we show below, it helps us reveal finer mechanisms and the role of cost of cohesion in driving the coevolutionary arms race dynamics. Here, we present an analytical approximation for this model based on the replicator dynamics and multilevel selection formalism. In S7 Appendix, we present a detailed derivation of the analytical model and compare the results with those from spatial simulations that explicitly include movement.

The population in this simplified model has four types of individuals: solitary cooperators, solitary defectors, cohesive cooperators and cohesive defectors. The dynamics of the system is completely defined by the relative frequencies of any three of these types, since the population size is held constant. The multilevel selection approach provides an intuitive and analytically tractable framework to model the dynamics of these different types: we consider the individual level selection pressure on cooperators separately within the cohesive and non-cohesive sub-populations, and the group level selection pressure between the cohesive and non-cohesive sub-populations. Since the cohesive and non-cohesive sub-populations are well-mixed, we can write replicator equations for the proportion of cooperators within each of them (denoted by *p*_*f*_ and *p*_*s*_, respectively). We can write another replicator equation for the proportion of cohesive individuals (*q*). The dynamics of the system is then described by the following differential equations (see S7 Appendix for a detailed analytical derivation):

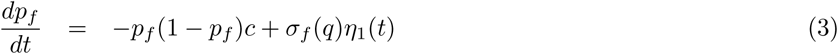

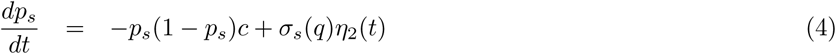

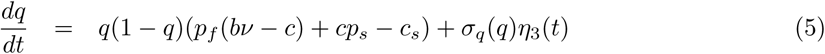

where *σ_i_* represents magnitude of noise due to stochastic group formation and mutations (described in detail in S7 Appendix), *η_i_*(*t*) are standard Gaussian white noise processes, and *ν* = (*n* - 1)/*n* with *n* being the average group size experienced by cohesive individuals. We also assume that the noise terms for the three equations are independent. Since the focus of this model is to obtain conditions for selection for cooperation and the arms race dynamics, we did not explicitly model mobility. We assumed that there is sufficient mobility to cause group formation among cohesive individuals. From the above three equations, we can show that dynamics of the overall proportion of cooperators in the population *p* can be written as

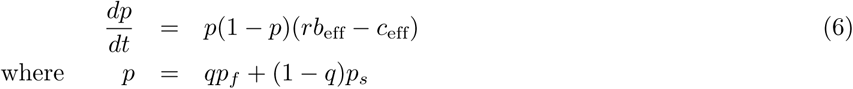

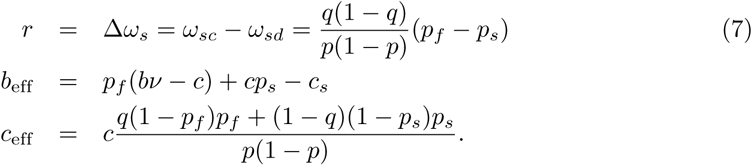

Here, if *b*_eff_(> -*c*_*s*_) and *c*_eff_(*>* 0) are treated as the effective benefits and costs incurred by cooperators, then *r* can be interpreted as the assortment coefficient (see S7 Appendix for derivation and more details). The effective benefit to cooperators (*b*_eff_) increases with the proportion of cohesive and solitary cooperators (*p*_*f*_ and *p*_*s*_), and is net positive only if the benefit from cooperation exceeds the cost of cohesion. The effective cost of cooperation (*c*_eff_) is equal to *c* when there is no variability of cooperators among cohesive and solitary types (*p*_*f*_ = *p*_*s*_), and reduces with increasing variability

#### Multilevel selection

Equations (3) and (4) describe ‘within-group’ selection against cooperation, and cause both cohesive cooperators (*p*_*f*_) and solitary cooperators (*p*_*s*_) to decrease. Even so, as equation (6) shows, cooperators can increase in the overall population if *rb*_eff_ *-c*_eff_ *>* 0. Biologically, the relation *r* = Δ*ω*_*s*_ implies that assortment is achieved by a differential cohesion between cooperators and defectors. Thus, our multilevel selection approach formally demonstrates our key simulation result from Fig 3C, and is consistent with the understanding of assortment developed using the inclusive fitness approach.

#### Arms race dynamics

Coupled stochastic differential equations (3-5) fully describe the coevolutionary dynamics of our system, but the noise terms in Equations (3-5) make it analytically intractable. Therefore we simulated these equations numerically using Ito interpretation. We find that, similar to our previous result of individual based models (Fig 3a), the average proportion of cooperators *p* (= *q p*_*f*_ + (1 - *q*) *p*_*s*_) and the average cohesive tendency *ω*_*s*_ (= *q*) do not reach a steady state (Fig 4a). All other results corresponding to the Figures 2 and 3 of the continuous trait model remain the same, in this simplified analytical model as well. However, the cyclical dynamics of Fig 3a become amplified, exhibiting four clearly distinguishable ‘phases’ in the time series of *ω*_*s*_ (Fig 4a): (a) Phase I, where the population has very low cohesive tendencies (*ω*_*s*_*≈* 0) with nearly all individuals being solitary (marked by magenta bands along the generations axis in Fig 4a), (b) Phase II, where the cohesion rapidly increases (blue bands), (c) Phase III with nearly all individuals being cohesive (*ω*_*s*_*≈* 1, grey bands), and (d) Phase IV, where the average cohesion reduces again (red bands). We explain this dynamic below.

**Fig 4.**
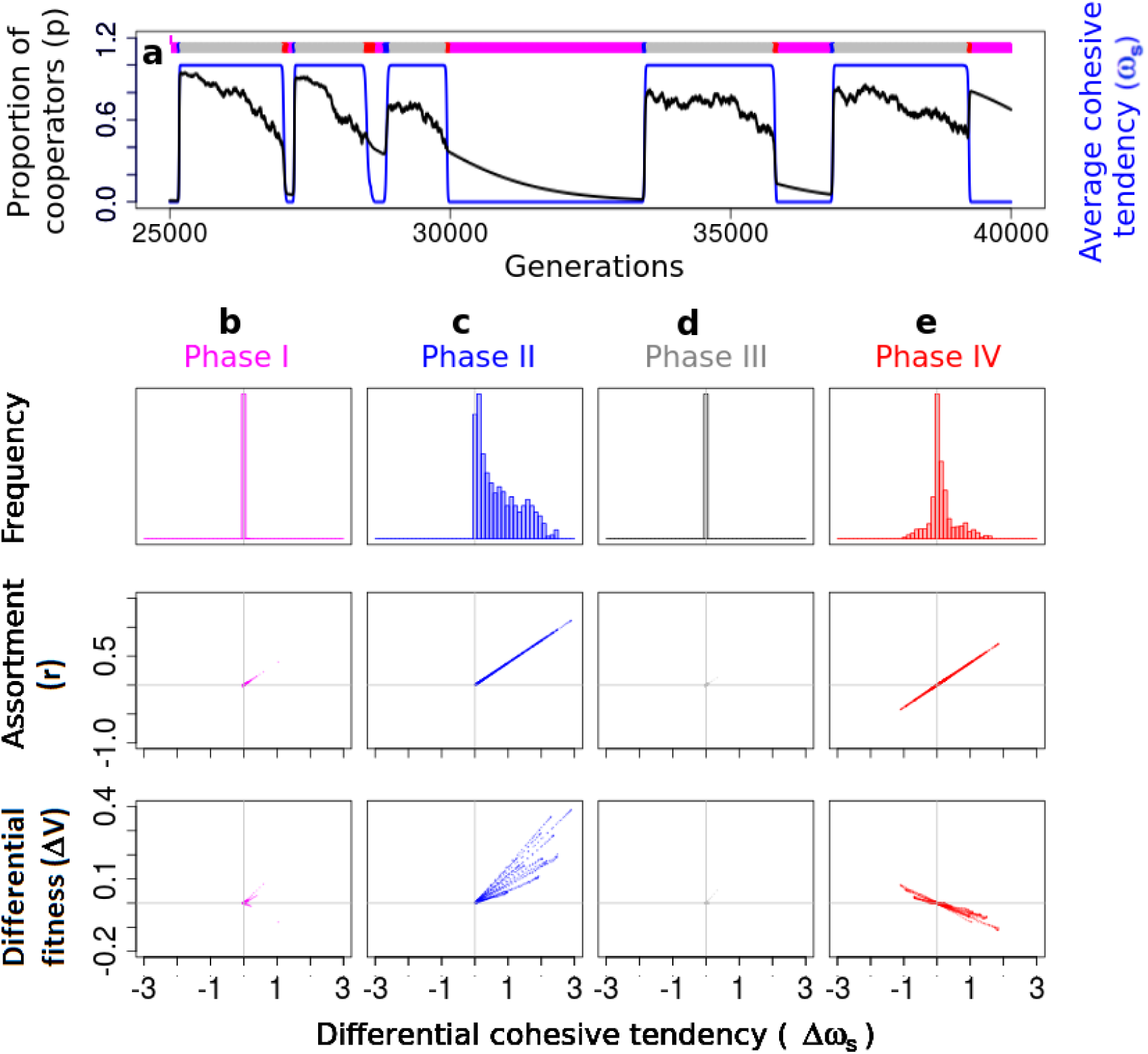
A simplified analytical model reveals cyclical arms race dynamics. (a) Analysis of a simpler analytical model reveals four phases of cyclical dynamics of cohesion (*ω*_*s*_; blue) and cooperation (*p*; black). The four phases are evident from the time series of *ω*_*s*_ marked by coloured bands along the generation axis. (b) Phase I (magenta): *ω*_*s*_ is close to zero with no differentialcohesion (Δ*ω*_*s*_ = *ω*_*sc*_ - *ω*_*sd*_ ≈ 0), thus no assortment (*r* ≈ 0) and no differential fitnes(Δ*V* = *V*_*c*_ - *V*_*d*_ ≈ 0) benefit to cooperators. (c) Phase II (blue): Cooperators lead an arms race to have a higher (costly) *ω*_*s*_ since they benefit by differential cohesion and assorting; but defectors benefit by matching *ω*_*s*_ of cooperators. (d) Phase III (grey): *ω*_*s*_ is high and same for all individuals, thus no assortment and no differential fitness benefit to cooperators. (e) Phase IV (red): Cooperators and defectors reduce their cohesive tendencies to avoid costs of cohesion and return to Phase I. The cyclical dynamics continues, as seen in (a). Parameters: See S7 Appendix

When the population consists of all non-cohesive individuals as in Phase I (*q* ≈ 0), the difference between cohesive tendencies of cooperators and defectors is close to zero (Δ*ω*_*s*_ ≈ 0; Fig 4b, d; top panel). In such a population which has a very few cohesive individuals, a few mutants that are both cooperative and cohesive cause *p*_*f*_ to sharply increase. As *p*_*f*_ rises beyond a threshold (*p*_*f*_ *> c*_*s*_*/*(*bν - c*)), groups of cohesive-cooperators form and get a higher pay-off than the rest of the population despite costs associated with both traits (Δ*V >* 0, Fig 4c; bottom panel). This initiates phase II, where we observe a rapid increase in the proportion of cohesive individuals (*dq/dt >* 0). This increase is driven by a consistent positive Δ*ω*_*s*_ (Fig 4c; top panel), akin to an arms race for the costly cohesive tendency. The above arms race continues until all individuals are cohesive (*q* ≈ 1), and thus the system reaches Phase III. Here, the differential cohesion and assortment is lost again (*r* = Δ*ω*_*s*_ ≈ 0;), putting cooperators at a disadvantage. As cooperators become rare (*p*_*f*_ *< c*_*s*_*/*(*bν - c*)), the benefits of grouping due to positive assortment can no longer offset the costs of cohesion. Thus, both cooperators and defectors now reduce their cohesive tendencies (*dq/dt <* 0) leading to Phase IV in which population quickly shifts to all solitary individuals. At the end of Phase IV, most individuals have no cohesive tendency, a scenario where cooperators are at a disadvantage in comparison to defectors. Therefore, the system is back to Phase I. The cycle then repeats. We represent the four phases on an *ω*_*s*_ vs *p* plot as in Fig 5. This figure also shows representative snapshots of how an equivalent spatial system would look, at various time points during these phases.

**Fig 5.**
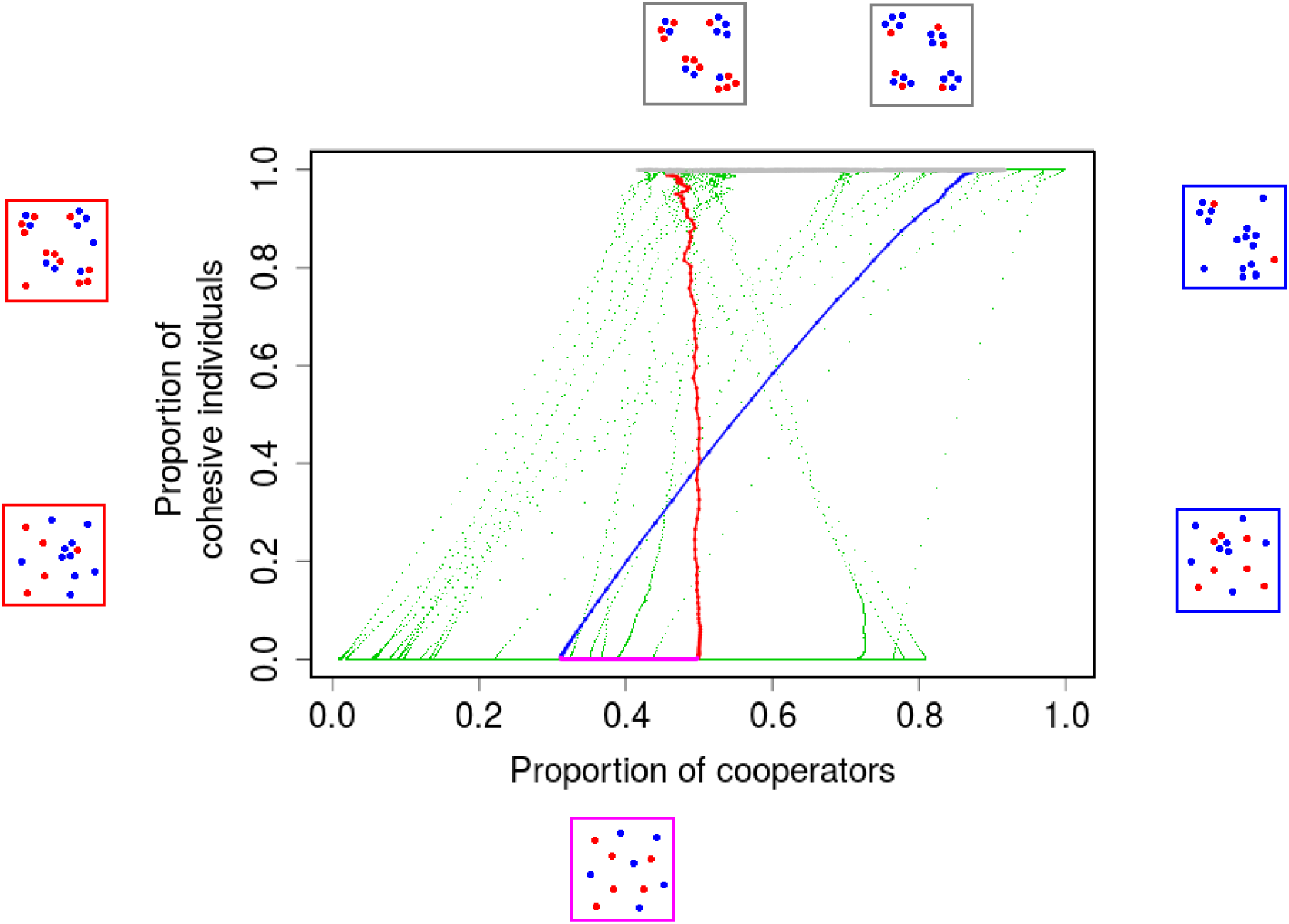
Cyclical coevolutionary dynamics of costly cooperative and cohesive traits along with schematic snapshots of expected spatial dynamics. (Spatial dynamics were not explicitly modelled analytially). Each data point (*p*,*ω*_*s*_) represents the instantaneous values of the proportion of cooperators (*p*) and average cohesive tendency (*ω*_*s*_) from the time series shown in Fig 4A. The four phases are evident here too and are coloured as in Fig 4 (For the sake of clarity, we have colour coded points from a single cycle, rest of the data points are in green). We show schematic snapshots representative of the expected group structure of the population, with cooperators in blue and defectors in red. In phase I, most individuals are solitary. Mutations lead to formation of cohesive and cooperative individuals and initiate the arms race to increase costly *ω*_*s*_, leading to Phase II which exhibits multilevel selection for cooperators. Nearly all individuals are cohesive in Phase III, destroying assortment and thus, reducing the level of cooperation. Finally, cooperators and defectors reduce the costly cohesive tendencies, thus reaching the Phase I back where *p* and *ω*_*s*_ are both low. These four phases repeat over generations.

Finally, we compare the timeseries obtained from the stochastic differential equations (SDEs) above and its analysis (Fig 4), with explicit spatial individual based simulations of individuals with a binary cohesive trait (S5 Figure). We find qualitative agreement between the two timeseries and their features.Further, S6 Figure shows the four phases from spatial simulations, which is in agreement with that obtained from the SDEs (Fig 5).

## Discussion

Our spatially explicit simulations demonstrate that mobility, arising from either self-propulsion or environmental forcing, can promote the evolution of cooperation via emergent group dynamics. Our models, including an analytical formulation, predict that cooperators and defectors can coexist by evolving differential strengths of cohesive interactions. This difference in local cohesion results in spontaneous assortment of cooperators. The cost of cohesion plays a crucial role in maintaining differential cohesive tendencies, thus preventing a run-away selection for cohesion that would eventually benefit defectors. Consequently, an arms race like dynamics ensues between cooperators and defectors for costly cohesive tendencies. Our proposed mechanism for the evolution of cooperation among non-kin depends on individual mobility and cohesive interactions. With these ingredients, we show that cooperation could occur even in the absence of limited dispersal or behavioural traits such as attraction to a phenotypic tag or moving away from unfavourable groups. Broadly, our study reveals that movement and emergent group dynamics, often thought to create mixing and thus destroy assortment, can facilitate coexistence of cooperation and defection among non-kin under a diverse range of ecological scenarios.

## Comparison with previous models and frameworks

Here, we discuss how the assumptions of our model preclude many known mechanisms of cooperation such as those arising from interactions between relatives, conditional strategies, and local dispersal. We then compare our results with greenbeard strategies and other studies employing an agent-based modelling framework and collective movement.

In our model we assumed that offspring disperse globally and move over relatively long time scales to reach an equilibrium group structure. Such large scale dispersal and mobility make the average relatedness of individuals within any local neighbourhood equal to the population average, as has been reported in fission-fusion societies [15]. This ruled out preferential interactions among genetic relatives within our model. Secondly, individuals could respond to nearby individuals’ positions and velocities (active system), or positions alone (passive system), but they had no ability to assess their cooperative behaviours, cohesive tendencies or the associated payoffs. Therefore, reciprocal strategies such as tit-for-tat [16] were ruled out. In addition, conditional mobility strategies such as inherent differential movement [47], ‘walk-away’ [20] or ‘success-driven migration’ strategies [48], where cooperators assess the neighbourhood and move away from non-cooperators and/or unfavourable environments were not applicable. Thirdly, studies have argued that spatial or network structures can facilitate assortment 49. These models typically assume a cost of dispersal and incorporate local dispersal of offspring that leads to spatial clustering of cooperators [7, 8, 10–12, 50–52]. By contrast, we assumed that there is no cost to dispersal and mobility, and that offspring disperse randomly in space after birth (global dispersal). Even if we include local dispersal in our model, initial assortment arising from local clustering is destroyed by the time the system reaches a fission-fusion group dynamic equilibrium (see S5 Appendix). If we include costs of dispersal and mobility within our model, we expect that results of our model may converge to the local dispersal driven clustering of cooperators [9 49].

Lehmann and Keller (2006) [53] classify grouping mechanisms as greenbeards. The idea of greenbeards was first proposed by Hamilton [3] and was later popularized by Dawkins [54, 55]. Greenbeards were originally defined as phenotypic traits that mediate interactions and are genetically linked to the cooperative trait [56, 57]. Over time, the concept of greenbeards has been generalized to any trait that generates assortment and is statistically linked to the cooperative trait [58]. By this definition, our differential cohesion mechanism is also a greenbeard. Theoretical studies have shown that greenbeard cooperation is prone to decay over time as defectors also evolve the greenbeard [59]However, recent studies [4, 5] have shown how greenbeard cooperation can arise in a cyclical manner as new tags are repeatedly discovered by cooperators, only to be subsequently invaded by defectors. Our model too shows cyclical behaviours of cooperation and defection (Fig 3), reminiscent of these studies. However, the mechanism described in these studies is unlikely to sustain cooperation when the tag under consideration is cohesion, because cooperators are always worse off than highly cohesive defectors. When cohesion is costly, cooperators can benefit by losing cohesion when they are rare. This offers them an escape route, allowing them to persist. Thus, our study highlights the role of the cost of cohesion in maintaining cooperation.

Our results complement the studies of Garcia, de Monte and colleagues who have highlighted the role of adhesion in the evolution of cooperation in mobile individuals with grouping tendencies 37, 45, 60. They showed that differential cohesion among mobile cooperators and defectors can lead to clustering of cooperators, resulting in fixation of cooperative trait in the population [37]. In the first two of the above studies, however, they assumed the existence of difference in adhesive interactions of cooperators and defectors. In the latter study, they allowed adhesion to evolve, but assumed that there is a correlation, or genetic linkage, between cooperative and adhesive traits. The above works also used public goods game where individuals gather some benefits of their own cooperative action. In our study, we used a modelling framework similar to theirs (agent based explicit spatial modelling). However, we assumed that the two traits (cooperation and cohesion/adhesion) could both evolve independently. Our main finding is that a correlation between these two traits, specifically a differential cohesion among cooperators and defectors, can arise spontaneously and is driven by an arms race dynamics for a costly cohesive trait. Indeed, if there are large differences between cooperators and defectors to begin with, our model predicts that the arms race dynamics acts to minimize those; however, cooperators will still maintain a slight edge in this dynamic. Moreover, we considered the more stringent case of costly cooperation by using prisoner’s dilemma game. We showed that differential adhesion based mechanism for the evolution of cooperation can emerge under stricter conditions than previously assumed.

Emergence of group-level sorting and information transfer is well studied in the context of mobile animal groups [35, 36]. For example, mobile groups can reach consensus decisions and/or arrive at foraging destinations even in the absence of any cues or signalling from, and abilities to recognise, individuals with pertinent information [61]; this is possible because mobility allows for separation of individuals with different movement properties (e.g. speed and cohesion) within and among groups [35, 38]. However, this principle has rarely been applied in the broader context of phenotypic assortment and its evolutionary implications. Our explicit incorporation of mobility showed that, counter to intuitive expectation, the fluidity caused by fission-fusion dynamics allows individuals to potentially interact with all other individuals, facilitating assortment of phenotypically similar individuals (cohesive cooperators) even when they are rare.

## Empirical implications

We now discuss the implications and potential applications of our model to real systems. We predict that cooperation can persist even in dynamic fission-fusion populations. Specifically, cooperation in organisms where motility and dispersal are not constrained by energetic or other costs could potentially be explained by our model. We note that mobility need not arise from self-propulsion alone; non-motile organisms may be driven, typically at no cost to individuals, by the dynamic environment (e.g. turbulent fluid) in which they are embedded (Fig 1). Such environmental conditions could be applicable to microorganisms living in turbulent oceanic environments [32], or to primitive unicellular organisms that may have lacked motility but lived in similar dynamic environments. Such systems could be employed to design experimental tests of our models. Other examples include microbial systems such as *Chlamydomonas*, which show aggregation correlated with size and motility [62], or microbes that exhibit an extreme form of cooperation requiring self-destruction, such as *Streptococcus pneumoniae* or *Salmonella typhimurium* which can infect the host only if the individuals producing the toxin lyse [30]. In these systems, local cohesion may arise from adhesiveness or stickiness between neighbouring individuals, a trait widely found in varying degrees across a range of microbial systems. It may also be worthwhile to investigate in such systems whether our proposed mechanism has a role to either initiate the evolution of cooperation or sustain a baseline level of cooperation, even if other mechanisms that support cooperation, such as phenotypic recognition or population viscosity, already exist.

Mobility can play a crucial role in pathogenesis of a number of diseases. A notable example is metastasis where tumour cells migrate from one organ to other. The process of migration may happen collectively as sheets or clusters of cells held together by adhesive molecules [63–65]. Moreover, these cells may switch between the different movement strategies depending on their micro-environmental conditions. Tumour cells are also thought to cooperate among themselves to overcome the body’s defences [66–68]. Understanding the evolutionary basis of metastasis may provide insights into developing strategies to prevent spread of the disease [65, 68]. Our simplified (binary-trait) model, which demonstrates cyclical dynamics and switching between collective and solitary movement along with the evolution of cooperation, can potentially offer insights into both proximate and ultimate factors driving the migration of tumour cells.

## Conclusion

In summary, we demonstrate that cooperation and fission-fusion dynamics can readily evolve together under a diverse range of ecological scenarios via multilevel-selection dynamics facilitated by self-sorting of groups. The self-sorted group structure itself emerges as a consequence of simple evolved local interactions among individuals even in the absence of limited dispersal, information on traits of other individuals, active group preferences, or dynamic cooperative strategies. The resulting macroscopic picture is that mobility, which is typically considered to inhibit the evolution of cooperation, can create conditions in which cooperation could, in fact, be much more prevalent. Our work suggests that collective movement could be an evolutionary adaptation that promotes cooperation over selfish interests among organisms.

## Supporting Information

**S1 Appendix Model and Implementation details**

**S2 Appendix Robustness to parameters**

**S3 Appendix Analytical calculations for simple cases using selection mutation equilibrium**

**S4 Appendix Deriving a measure of assortment**

**S5 Appendix Effect of model variations**

**S6 Appendix Passive particles system**

**S7 Appendix Analytical calculations of the simplified binary-trait model**

**S1 Video. Movement types** Representative videos showing the range of movement types that our model can exhibit, by changing the cohesive tendency of individuals. (Cohesive tendency is *R*_*s*_ in the active case and *γ* in the passive case; see S1 Appendix). In the passive system case, the background colours show the potential field of the turbulent fluid. For very low cohesive tendencies, individuals remain mostly solitary. For intermediate values of cohesive tendency, we get fission-fusion groups. For large values of cohesive tendency, we get very few large groups. In these simulations, the cohesive tendency of all individuals is identical.

**S2 Video. Self Sorting** Representative videos showing assortment due to a difference in cohesive tendencies of cooperators and defectors. There is no assortment when the cohesive tendencies (*R*_*s*_ in passive case and *γ* in active case) of cooperators and defectors are same. In these simulations, we have exaggerated the difference to make the assortment visually apparent.

**S1 Figure Robustness of results to model parameters.**

**S2 Figure Robustness of results to model variations.**

**S3 Figure Coevolutionary dynamics results for the passive system.**

**S4 Figure Replicator dynamics without noise.**

**S5 Figure Arms race and coevolutionary dynamics with explicit spatial simulations.**

**S6 Figure Phase portrait with explicit spatial simulations.**

## Acknowledgments

We thank Carey Nadell, Vidyanand Nanjundiah and Corina Tarnita for critical discussion and comments on this work. We also thank two anonymous reviewers for extremely detailed and insightful comments on the manuscript.

